# Improvements in the fossil record may largely resolve the conflict between morphological and molecular estimates of mammal phylogeny

**DOI:** 10.1101/373191

**Authors:** Robin M. D. Beck, Charles Baillie

**Affiliations:** School of Environment and Life Sciences, University of Salford, Manchester M5 4WT

## Abstract

Morphological phylogenies of mammals continue to show major conflicts with the robust molecular consensus view of their relationships. This raises doubts as to whether current morphological character sets are able to accurately resolve mammal relationships, particularly for fossil taxa for which, in most cases, molecular data is unlikely to ever become available. We tested this under a hypothetical “best case scenario” by using ancestral state reconstruction (under both maximum parsimony and maximum likelihood) to infer the morphologies of fossil ancestors for all clades present in a recent comprehensive molecular phylogeny of mammals, and then seeing what effect inclusion of these predicted ancestors had on unconstrained analyses of morphological data. We found that this resulted in topologies that are highly congruent with the molecular consensus, even when simulating the effect of incomplete fossilisation. Most strikingly, several analyses recovered monophyly of clades that have never been found in previous morphology-only studies, such as Afrotheria and Laurasiatheria. Our results suggest that, at least in principle, improvements in the fossil record may be sufficient to largely reconcile morphological and molecular phylogenies of mammals, even with current morphological character sets.

## Introduction

The evolutionary relationships of mammals have been a major focus of research within systematics for over a century [1-6]. In the last two decades, the increasing availability of molecular data has seen the emergence of a robust “consensus” phylogeny of extant mammals This consensus indicates that several groupings of placental mammals proposed based on morphological data (such as “edentates”, “ungulates”, and “insectivorans”) are polyphyletic [4-6]. It also shows that living placentals are distributed among four major clades or “superorders”, likely reflecting their biogeographical history, namely Xenarthra, Afrotheria, Laurasiatheria and Euarchontoglires [4-6]. Arguably the most striking finding is that both Afrotheria and Laurasiatheria include “insectivoran-grade” (afrotherian tenrecs and golden moles, laurasiatherian eulipotyphlans such as hedgehogs, shrews and moles), “ungulate-grade” (afrotherian proboscideans, hyraxes and sea cows, laurasiatherian artiodactyls and perissodactyls), and myrmecophagous (afrotherian aardvarks, laurasiatherian pangolins) members. There is strong molecular evidence that Laurasiatheria and Euarchontoglires are sister-taxa, forming Boreoeutheria [4-6], and it seems probable that Xenarthra and Afrotheria also form a clade, named Atlantogenata [4, 7, 8] (Figure 1).

**Figure 1.**
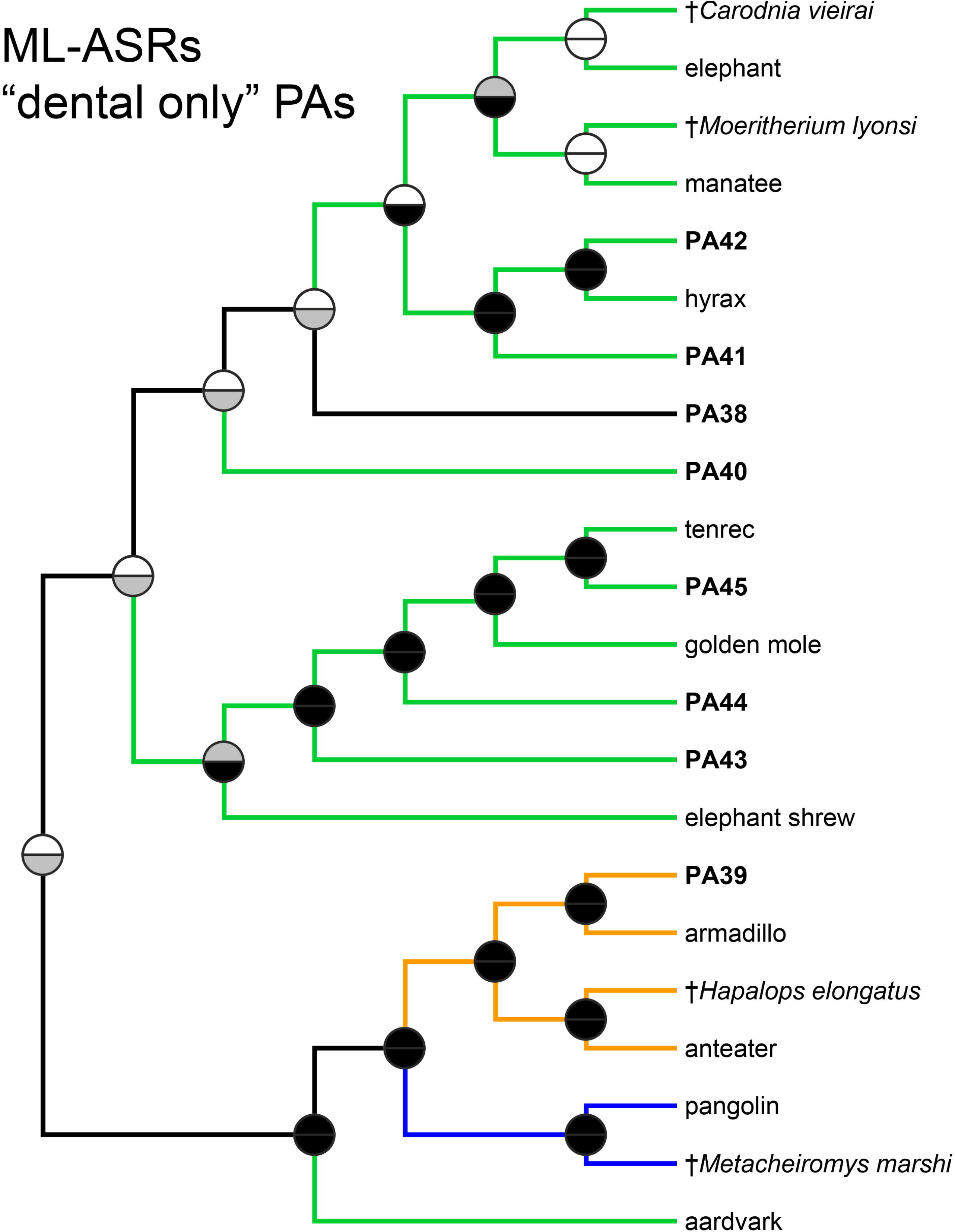
Molecular (a) and morphological (b) phylogenies of mammals. The topology shown in (a) is modified from Fig. 1 of Meredith et al. and is based on an 11,000 amino acid alignment from 26 gene fragments, whereas that in (b) is modified from Figure S2A of O’Leary et al. and is based on 4541 morphological characters (fossil taxa have been deleted). Branches are colour coded according to their membership of the superorders Afrotheria, Xenarthra, Laurasiatheria and Euarchontoglires. Node numbers in (a) correspond to the predicted ancestors for those nodes in Figure 2.

Recent morphology-only analyses of mammal relationships continue to show numerous areas of conflict with this molecular consensus, and typically recover “insectivoran”, “ungulate“, and “edentate” clades [3, 9-11] (Figure 1). “Total evidence” analyses that combine morphological and molecular data are largely congruent with the molecular consensus [3, 9], but “pseudoextinction” analyses of total evidence datasets, in which selected extant placentals are treated as if they are fossils by deleting their molecular data, often fail to recover the pseudoextinct taxa within their respective superorders [12]. This has led to doubts regarding the ability of morphological data to accurately reconstruct the evolutionary relationships of mammals [10, 12-14]. In particular, it raises questions as to whether genuine fossil taxa (for most of which molecular data is unlikely to ever become available) can be correctly placed within mammalian phylogeny, even when using a total evidence approach. This creates a dilemma, because the inclusion of fossil taxa may be critically important for phylogenetic comparative analyses [15-18], and yet such analyses assume that the fossils are placed accurately in the phylogeny being used.

In general, we should expect increased taxon sampling to improve phylogenetic accuracy [19-21]. In morphological and total evidence analyses, fossil taxa are likely to be particularly important: by exhibiting unique combinations of plesiomorphic and derived character states, they should help break up long morphological branches leading to extant taxa [20, 22, 23]. The most detailed morphological study of mammal phylogeny published to date is that of O’Leary et al. [3], but this is still highly incongruent with the molecular consensus [10] (Fig. 1). However, O’Leary et al. [3] included only 40 fossil taxa, which is a tiny fraction of the number likely to have existed during the Mesozoic and Cenozoic [24]. A key question then, is whether improving the taxon sampling of morphological analyses, particularly of fossil taxa, might be sufficient to resolve the conflict between morphological and molecular estimates of mammal phylogeny.

We investigate this by first reconstructing the expected morphological character states (based on the O’Leary et al. [3] matrix) of the fossil ancestors of all of the clades present in the molecular consensus, and then testing what impact the inclusion of these predicted fossil ancestors has on the results of morphology-only analyses. In effect, our analyses represent a hypothetical “best case scenario” for morphological studies of mammal phylogeny, in which we simulate the discovery of direct fossil ancestors.

If the inclusion of these predicted ancestors in morphology-only analyses is sufficient to result in phylogenies that are largely congruent with the molecular consensus, then it suggests that the current conflict between morphological and molecular estimates of mammal phylogeny might be resolved by improvements in the fossil record and the addition of more fossil taxa to currently available morphological matrices. In turn, this would suggest that fossil taxa (including non-ancestral forms), for which molecular data is unavailable, could still be accurately placed within mammal phylogeny given sufficiently dense taxon sampling, and hence that phylogenies that include fossil and extant mammals may become sufficiently accurate for use in comparative analyses. Conversely, if morphology-only analyses remain strongly incongruent with the molecular consensus, even under this hypothetical “best case scenario”, then it suggests that the conflict will not be resolved simply by improved taxon sampling, and that the relationships of fossil taxa inferred using current datasets and methods of analysis should not be viewed with confidence.

## Methods

The morphological matrix employed here is the version used by O’Leary et al. [3] for their model-based analyses, comprising 4541 characters scored for 46 extant and 40 fossil taxa. We modified the matrix by first merging the character scores for the fossil notoungulate *Thomashuxleya externa* with those from a more recent study [25], and then deleting 407 constant characters. This left a total of 4134 characters: 1170 cranial, 1311 dental, 890 postcranial, and 763 soft tissue. For the ancestral state reconstructions (ASRs), all fossil taxa were deleted from the matrix, and we assumed the molecular topology present in Fig. 1 of Meredith et al. [4] (Figure 1a). We used two different optimality criteria for inferring ASRs: maximum parsimony (MP) and maximum likelihood (ML). MP-ASRs for all nodes were calculated using the “Trace All Characters” command in *Mesquite*, whereas ML-ASRs were calculated in *RAxML* using the “-f A” command (which calculates marginal ancestral states), assuming the MK+GAMMA model, and applying the Lewis correction (absence of invariant sites) for ascertainment bias [26]. The collective MP-ASRs for each node were then added to the original matrix (i.e. with all extant and fossil taxa present), to act as predicted ancestors (PAs). The same was done for the ML-ASRs, resulting in two different matrices: one with PAs based on MP-ASRs, and one with PAs based on ML-ASRs.

Different anatomical partitions differ in their preservation probability: in general, soft tissue characters are the least likely to preserve, followed by postcranial, then cranial, and lastly dental characters [27, 28]. We further modified these three matrices to take this into account: in the “all characters” version of the matrix, we retained all character scores for the PAs; for “skeletal only” we scored all soft tissue characters as unknown for the PAs; for “craniodental only” we scored all soft tissue and postcranial characters as unknown for the PAs; for “dental only”, we only included dental character scores for the PAs. To further investigate the impact of fossil preservation, we used a custom *R* script to only retain character scores for the PAs that could be scored in: 1) at least one of the 40 “real” fossil taxa in the matrix, simulating a scenario in which the PAs are extremely well-preserved (= “max preservation”); or 2) in at least 50% (i.e. at least 20) of the “real” fossil taxa, simulating a scenario in which the PAs show a “typical” or “average” degree of preservation (= “typical preservation”). All matrices are available in the electronic supplementary material (Data File S1).

For the MP-ASRs, the original matrix and the different versions of the matrix with PAs included were analysed using MP, as implemented by *TNT*. Tree searches comprised new technology searches until the same minimum length was hit 100 times, followed by a traditional search with TBR branch-swapping among the trees already saved. All most parsimonious trees were summarised using strict consensus, and bootstrap support values were calculated as absolute frequencies using 500 replicates. For the ML-ASRs, the original matrix and the different versions of the matrix with PAs included were analysed using *RAxML*, again with the MK+GAMMA model and the Lewis correction for ascertainment bias. *RAxML* searches comprised 1000 replicates of the default rapid hill-climbing algorithm. Non-parametric bootstrap support values were also calculated, with the number of replicates determined by the autoMRE criterion. Standard bootstrap values may be unduly conservative when one or more “rogue” taxa are present, and so we also calculated support for all our MP and ML trees using the recently-developed “transfer bootstrap expectation” (TBE) method [29], using *BOOSTER*. All trees and associated support values are available in the electronic supplementary material (Data File S2).

After deleting all fossil taxa and PAs, the trees from all analyses were quantitatively compared with the same Meredith et al. [4] topology used to calculate the ASRs. We used the normalised Robinson-Foulds (= partition) metric (nRF), the SPR distance (SPRd), and the distortion coefficient (DC); the latter two metrics are less affected by shifts in position of just one or a few taxa [30]. A greater degree of similarity between trees is indicated by values closer to 0 for nRF, but values closer to 1 for SPRd and DC. In the case of MP analyses that recovered more than one most parsimonious trees, we compared the individual most parsimonious trees, and also the strict consensus of these, to the Meredith et al. [4] topology.

## Results and discussion

Strikingly, for many analyses, inclusion of PAs was sufficient to result in phylogenies that are generally congruent with the molecular consensus, particularly those that assumed better preserved PAs (Table 1; Figure 2). For example, for the MP analysis assuming “maximum fossilisation” PAs, the strict consensus recovers monophyly of Atlantogenata, Boreoeutheria, and all four superorders (Figure 2). Even when strict monophyly of the four superorders was not recovered, this was often due to only one or a few taxa being misplaced, as indicated by high SPRd and DC values (Table 1). Where monophyly of Afrotheria and Laurasiatheria was not recovered, this was often due to misplacement of the aardvark and the pangolin (Figure 3; electronic supplementary material, Data File S2); these two taxa are characterised by a greatly simplified (aardvark) or entirely absent (pangolin) dentition, and so cannot be scored for many dental characters (~32% of the characters in the original morphological matrix of O’Leary et al. [3] are dental).

**Table 1.** Summary of results of phylogenetic analyses with and without predicted ancestors (PAs) under maximum parsimony (MP) and maximum likelihood (ML). Fit relative to the molecular phylogeny of Meredith et al. was assessed using the normalised Robinson-Foulds metric (nRF), the SPR distance (SPRd), and the distortion coefficient (DC). nRF values closer to 0 and SPRd and DC values closer to 1 represent better fit. For the MP analyses that recovered more than one most parsimonious tree (MPT), the number of MPTs is indicated. Also indicated is whether the analyses recovered six superordinal clades found in the “molecular consensus”, with standard bootstrap (BS) and “transfer bootstrap expectation” (TBE) values given in brackets. ^a^Values vary between MPTs – value given here represents the mean value weighted by the number of MPTs

| Analysis | nRF | SPRd | DC | Monophyletic? |  |  |  |  |  |
| --- | --- | --- | --- | --- | --- | --- | --- | --- | --- |
|  |  |  |  | Atlantogenata | Afrotheria | Xenarthra | Boreoeutheria | Euarchontoglires | Laurasiatheria |
| **MP** |  |  |  |  |  |  |  |  |  |
| **no PAs** | 0.57 | 0.56 | 0.70 | no | no | **yes** (BS = 74;<br>TBE = 87) | no | no | no |
| **“all characters” PAs** |  |  |  |  |  |  |  |  |  |
| MPTs (n = 64) | 0.23 | 0.86 | 0.94 | **yes** (BS < 50;<br>TBE < 50) | **yes** (BS < 50;<br>TBE < 50) | **yes** (BS = 63;<br>TBE = 75) | **yes** (BS < 50;<br>TBE < 50) | **yes** (BS = 90; TBE<br>= 95) | **yes** (BS < 50;<br>TBE < 50) |
| strict consensus | 0.24 | 0.88 | 0.93 |  |  |  |  |  |  |
| **“skeletal only” PAs** |  |  |  |  |  |  |  |  |  |
| MPTs (n = 64) | 0.23 | 0.86 | 0.94 | **yes** (BS < 50;<br>TBE < 50) | **yes** (BS < 50;<br>TBE < 50) | **yes** (BS = 64;<br>TBE = 76) | **yes** (BS < 50;<br>TBE < 50) | **yes** (BS = 80; TBE<br>= 89) | **yes** (BS < 50;<br>TBE < 50) |
| strict consensus | 0.24 | 0.88 | 0.93 |  |  |  |  |  |  |
| **“craniodental only” PAs** |  |  |  |  |  |  |  |  |  |
| MPTs (n = 12) | 0.46 | 0.70 | 0.85 | no | no | **yes** (BS < 50;<br>TBE = 65) | no | **yes** (BS = 59; TBE<br>= 78) | no |
| strict consensus | 0.47 | 0.70 | 0.84 |  |  |  |  |  |  |
| **“dental only” PAs** |  |  |  |  |  |  |  |  |  |
| MPTs (n = 30) | 0.54 | 0.63<sup>a</sup> | 0.76<sup>a</sup> | no | no | **yes** (BS < 50;<br>TBE = 58) | no | **yes** (BS < 50; TBE<br>< 50) | no |
| strict consensus | 0.46 | 0.84 | 0.65 |  |  |  |  |  |  |
| **“max preservation” PAs** |  |  |  |  |  |  |  |  |  |
| MPTs (n = 64) | 0.23 | 0.86 | 0.94 | **yes** (BS < 50;<br>TBE < 50) | **yes** (BS < 50;<br>TBE < 50) | **yes** (BS = 63;<br>TBE = 75) | **yes** (BS < 50;<br>TBE < 50) | **yes** (BS = 81; TBE<br>= 90) | **yes** (BS < 50;<br>TBE < 50) |
| strict consensus | 0.24 | 0.88 | 0.93 |  |  |  |  |  |  |
| **“typical preservation” PAs** |  |  |  |  |  |  |  |  |  |
| MPTs (n = 2) | 0.46 | 0.72 | 0.86 | no | no | **yes** (BS < 50;<br>TBE = 65) | no | **yes** (BS < 50; TBE<br>= 62) | no |
| strict consensus | 0.48 | 0.72 | 0.85 |  |  |  |  |  |  |
| **ML** |  |  |  |  |  |  |  |  |  |
| **no PAs** | 0.59 | 0.56 | 0.70 | no | no | **yes** (BS = 96;<br>TBE = 98) | no | no | no |
| **“all characters” PAs** | 0.21 | 0.81 | 0.95 | **yes** (BS < 50;<br>TBE = 94) | no | **yes** (BS = 91;<br>TBE = 94) | **yes** (BS < 50;<br>TBE = 79) | **yes** (BS = 56; TBE<br>= 79) | **yes** (BS < 50;<br>TBE = 91) |
| **“skeletal only” PAs** | 0.25 | 0.81 | 0.94 | **yes** (BS < 50;<br>TBE = 93) | no | **yes** (BS = 93;<br>TBE = 95) | no | no | **yes** (BS < 50;<br>TBE = 88) |
| **“craniodental only” PAs** | 0.32 | 0.81 | 0.91 | no | **yes** (BS < 50;<br>TBE = 65) | **yes** (BS = 88;<br>TBE = 92) | no | no | no |
| **“dental only” PAs** | 0.41 | 0.74 | 0.88 | no | no | **yes** (BS = 97;<br>TBE = 98) | no | no | no |
| **“max preservation” PAs** | 0.27 | 0.81 | 0.93 | no | **yes** (BS = 91;<br>TBE = 94) | no | no | **yes** (BS < 50; TBE<br>= 89) | no |
| **“typical preservation” PAs** | 0.43 | 0.74 | 0.87 | no | no | **yes** (BS = 76;<br>TBE = 84) | no | no | no |

**Figure 2.**
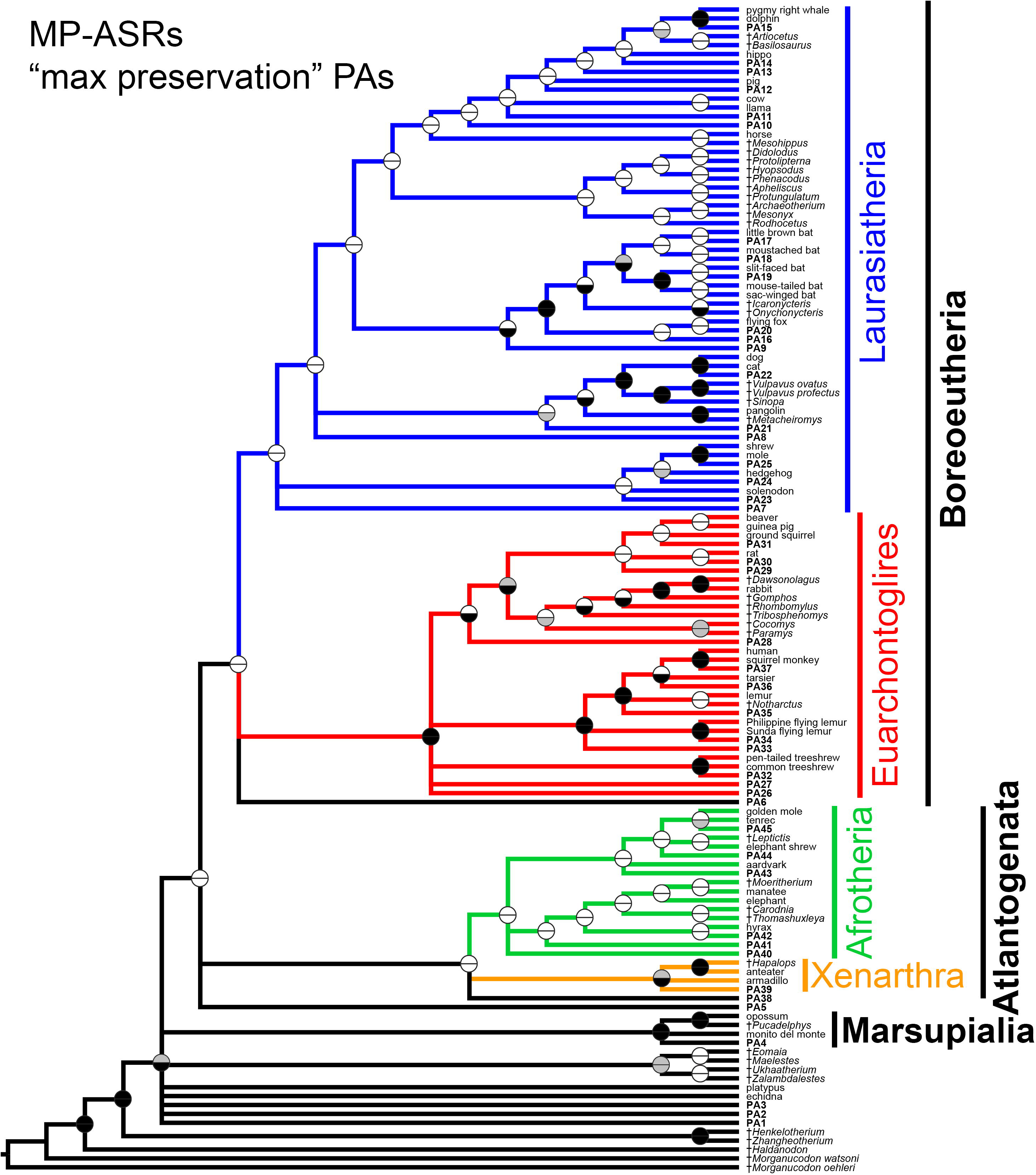
Morphological phylogeny of mammals based on maximum parsimony (MP) analysis of a modified version of the dataset of O’Leary et al. and with “max preservation” predicted ancestors (PAs) added. Topology shown is a strict consensus of 64 most parsimonious trees. Ancestral states for the PAs were reconstructed using MP. PAs are shown in bold, with numbers corresponding to the nodes for which they are ancestral, as shown in Figure 1a. Fossil taxa are indicated with †. Circles at nodes indicate support, with the top half representing standard bootstrap values and the bottom half “transfer bootstrap expectation” (TBE) values: black indicates ≥70% support, grey 50-69%, and white <50%.

Although a major source of phylogenetic information, dental characters have been argued to be less reliable for recovering mammalian phylogeny than are characters from other parts of the skeleton [27]. However, when PAs were represented by dental characters only, ML analysis recovered a clade that includes both “insectivoran-grade” and “ungulate-grade” afrotherians (the aardvark was placed in an “edentate” clade that also includes the pangolin, the fossil pangolin relative *Metacheiromys*, and xenarthrans); this clade receives moderate (65%) TBE support (Figure 3). This analysis also grouped “insectivoran-grade” and “ungulate-grade” laurasiatherians together with bats, with high (83%) TBE support (electronic supplementary material, Data File S2); however, carnivorans and the pangolin were absent from this clade, and so true laurasiatherian monophyly was not supported (electronic supplementary material, Data File S2). Neverthless, these results suggest that the discovery and inclusion of ancestral fossils known only from teeth may be sufficient to greatly improve the congruence between morphological and molecular phylogenies of mammals.

**Figure 3.**
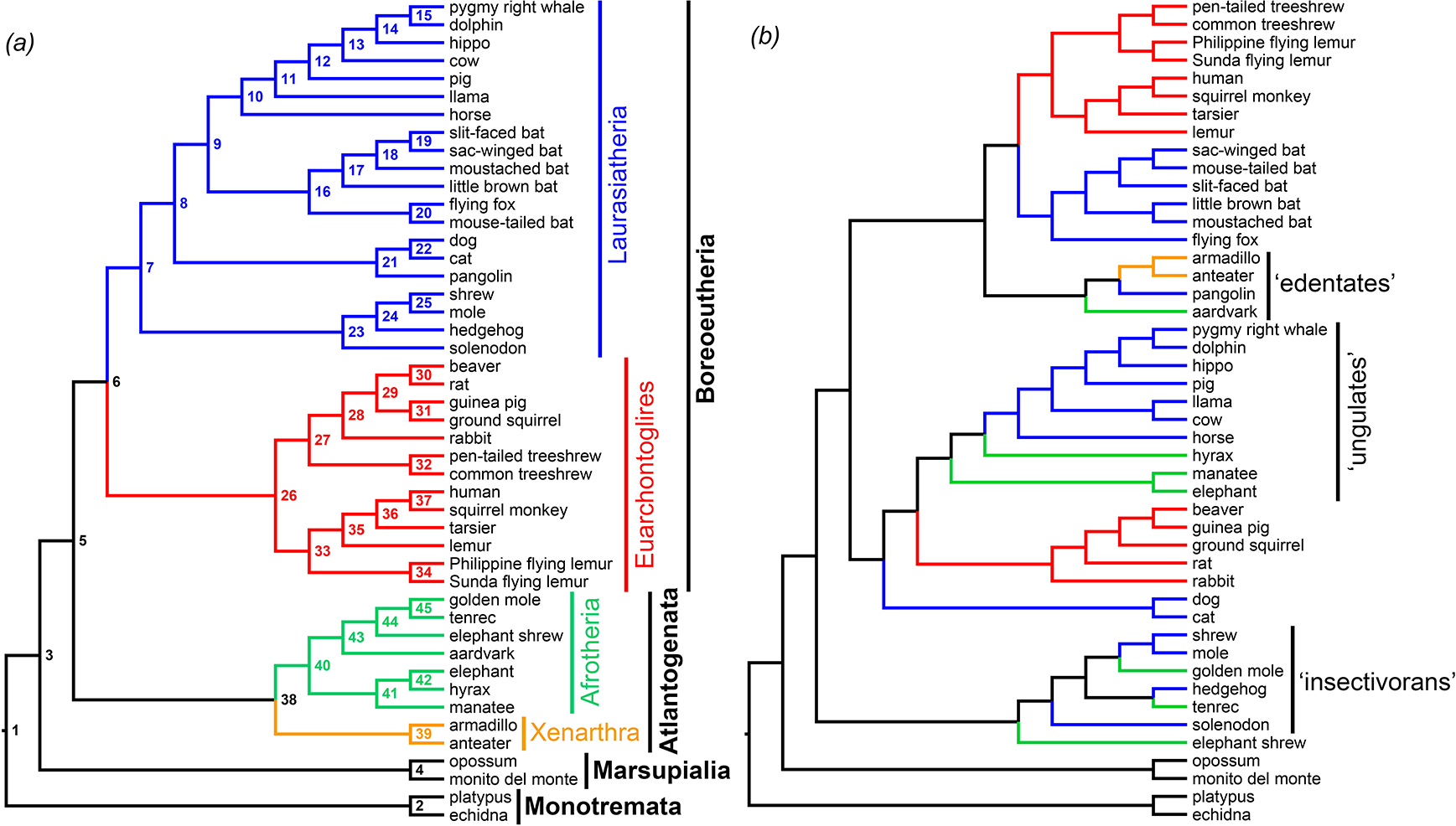
Partial morphological phylogeny of mammals based on maximum likelihood (ML) analysis of a modified version of the dataset of O’Leary et al. and with “dental only” predicted ancestors (PAs) added; only the part of the tree that includes afrotherians is shown here. Ancestral states for the PAs were reconstructed using ML. PAs are shown in bold, with numbers corresponding to the nodes for which they are ancestral, as shown in Figure 1a. Fossil taxa are indicated with †. Circles at nodes indicate support, with the top half representing standard bootstrap values and the bottom half “transfer bootstrap expectation” (TBE) values: black indicates ≥70% support, grey 50-69%, and white <50%.

Bootstrap values are low for most nodes in the MP analyses, even when using the TBE method (Figure 2; electronic supplementary material, Data File S2), but this might be expected given the inclusion of multiple fossil taxa (both “real” fossil taxa and PAs) [23, 31]. Support values were generally higher for the ML analyses, particularly when using the TBE method (Figure 3; electronic supplementary material, Data File S2), including for several of the molecular consensus clades (where recovered; Table 1; electronic supplementary material, Data File S2).

In general, the MP analyses were somewhat more successful in recovering the clades that characterise molecular consensus than were the ML analyses. One possible explanation for this is that the ML ancestral state reconstruction assumed a single set of branch lengths for all characters, which is likely to be inappropriate for morphology [30, 32]; future studies could investigate the impact of data partitioning [33]. Another obvious area to explore would be the use of tip-dating approaches [34-36], with PAs assigned ages compatible with recent molecular clock analyses (e.g. [4]); recent work suggests that such approaches may be better able to identify cases of homoplastic resemblance than methods that do not incorporate temporal evidence [37].

We emphasise that our study represents a hypothetical “best case scenario”: we inferred the ancestral states of PAs using the same approach (either MP or ML using the Mkv+G model) that was subsequently used to analyse the matrix with the PAs added; the PAs lack any apomorphies not present in their descendants (i.e. they represent “perfect” ancestral morphologies); and we included a PA for every single clade present in the Meredith et al. [4] tree. All of these are unrealistic, or at least highly optimistic, assumptions (although direct ancestors may actually be relatively common in the fossil record; [38, 39]). We also did not assess whether the character combinations of the PAs appear biologically plausible or not.

Nevertheless, we show that the inclusion of PAs predicted by the molecular consensus of placental phylogeny is sufficient to result in morphological phylogenies that closely match this consensus, without the use of constraints or the addition of molecular data, even when incomplete fossilisation is taken into account, and that there are at least hypothetical character combinations that can link morphologically disparate mammalian taxa, such as the “insectivoran-grade”, “ungulate-grade” and myrmecophagous members of Afrotheria and Laurasiatheria. If genuine fossil taxa exhibit these character combinations, then their discovery and inclusion in phylogenetic analyses might be sufficient to largely resolve the current conflict between molecular and morphological analyses, even using morphological matrices that currently show extensive conflict with molecular data, such as that of O’Leary et al. [3]. There is increasing evidence that this may indeed be the case; for example, well-preserved remains of *Ocepeia* from the Palaeocene of Morocco reveal that it combines features of “insectivoran” and “ungulate” afrotherians [40], and other African fossils show that dental similarities between “ungulate” afrotherians and laurasiatherians are homoplastic [41].

Although not tested here, a similar principle may apply to other clades, if so, then we can be optimistic that we may be able to accurately infer phylogeny for many clades using morphological data alone, given a sufficiently good fossil record. Improvements in phylogenetic methods will undoubtedly also play a role: these might include better models of morphological character evolution [42, 43], clock models [34-36], methods that take into account character non-independence and saturation [44, 45], or some combination of these. However, our results suggest that inclusion of fossil taxa may prove to be particularly important. In any case, arguments that morphological data is “inadequate” for accurately inferring the phylogeny of mammals [12-14], or of the many other clades that currently show extensive morphological-molecular conflict (such as birds [46-49]), are at the very least premature.

## Acknowledgements

We thank Marcelo Marcelo R. Sánchez-Villagra (University of Zürich) and Robert Asher (University of Cambridge) for discussion.

